# Splice-switching antisense oligonucleotides correct cryptic exon inclusion and restore SDCCAG8 protein in Bardet-Biedl Syndrome

**DOI:** 10.1101/2025.10.15.682674

**Authors:** Kelleen E McEntee, Bailey L McCurdy, Austin Larson, Emily A McCourt, Michael L Kaufman, Amy E Campbell, Chad G Pearson, Scott Demarest, J Matthew Taliaferro, Jay R Hesselberth, Sujatha Jagannathan

## Abstract

Bardet-Biedl Syndrome (BBS) is a ciliopathy often associated with progressive blindness and obesity. A patient presenting with BBS was discovered to have two mutations within 55bp of each other in intron 7 of *SDCCAG8* (*BBS16*). One of the biallelic mutations, c.740+356C>T, causes inclusion of cryptic exon(s) containing premature termination codons, while c.740+301G>A has not been characterized. We hypothesized that antisense oligonucleotides (ASOs) complementary to the patient’s mutations or to the cryptic exon splice sites would correct the splicing of *SDCCAG8* between exons 7 and 8 to prevent cryptic exon inclusion and restore *SDCCAG8* expression. We systematically screened 20nt-long ASOs tiled across each mutation and ASOs targeting the 3′ splice sites of the cryptic exons in patient-derived fibroblasts, using RT-PCR assays to assess exon 7 and 8 splicing. We identified one ASO for each mutation and a cryptic exon-targeting ASO that restored the splicing pattern to that observed in an unaffected cell line. Lead ASOs were further investigated through RT-PCR, RNA sequencing, and western blotting to confirm ASO-mediated restoration of wild-type transcript and protein. Notably, ASO 20, which targets the cryptic exon 7a/7a′ splice site rather than patient-specific mutations, achieved the greatest rescue effect, increasing exon 7-8 splicing from 0% to an average of 26% and restoring SDCCAG8 protein from undetectable levels to approximately 40% of wild-type expression. This mutation-agnostic approach could benefit multiple patients with cryptic exon inclusion in this region of *SDCCAG8*, expanding therapeutic impact beyond traditional N-of-1 ASO strategies. These findings establish a molecular foundation for clinical development of ASO therapy for BBS caused by *SDCCAG8* splicing defects.

## INTRODUCTION

Antisense oligonucleotides (ASOs) are short oligonucleotides that can be tailored to treat specific diseases and mutations. ASOs bind to transcripts and can alter their splicing patterns to treat diseases such as Dravet syndrome and Spinal muscular atrophy [1,2]. The mechanism of these therapies depends on the mutation and its impact on gene expression. However, most function by masking sequence elements in pre-mRNA that would otherwise result in the degradation of the transcript or production of truncated proteins [3]. Restoration of correct splicing and full-length transcript is therefore the primary therapeutic goal, as it enables production of functional protein. N-of-1 ASOs, i.e., ASOs designed to treat one individual and target their specific mutation, have been used to treat individuals within a year of drug development, such as Milasen [4], highlighting the potential of this approach [3]. Bardet-Biedl Syndrome (BBS) is a rare autosomal recessive ciliopathy that results in retinal degeneration and other symptoms [5]. BBS is caused by the malfunction or absence of proteins that disrupt primary cilia formation and/or function [5]. Serologically defined colon cancer autoantigen protein, SDCCAG8 (also referred to as BBS16, NPHP10, and CCCAP), is a protein that localizes to the centrosome and basal body where it plays critical roles in ciliogenesis and hedgehog signaling [6–8] . Mutations in *SDCCAG8* that disrupt these functions can result in BBS and other ciliopathies [9].

A patient diagnosed with BBS was found to have two mutations in *SDCCAG8*, c.740+301G>A and c.740+356C>T located within 55 bp of each other in intron 7. One of these mutations, c.740+356C>T, interferes with splicing and is thought to cause the loss of an exonic splicing enhancer that promotes the inclusion of cryptic exon(s) which introduces a premature termination codon (PTC) [7]. c.740+301G>A is classified as a variant of unknown significance (VUS) in ClinVar and is predicted to create a 5′ splice donor [10]. The presence of a PTC may cause the degradation of the transcript via nonsense-mediated RNA decay (NMD) or the production of a truncated protein lacking critical functional domains [11]. In the case of SDCCAG8, loss of the C-terminal domain is expected to eliminate centrosome localization, thereby abolishing protein function [12,13].

Currently, there is no treatment that prevents or slows BBS-related retinal degeneration, and existing treatments only target the symptoms of BBS [5]. We hypothesized that masking the patient’s mutations or the cryptic splice sites with ASOs would prevent inclusion of cryptic exons and restore full-length *SDCCAG8* transcript. The resulting restoration of full-length SDCCAG8 protein would preserve the C-terminal domain required for centrosome localization, potentially restoring protein function.

## MATERIALS AND METHODS

### Cell culture

Patient-derived and unaffected human dermal fibroblasts (106-05A; Sigma-Aldrich) were cultured in MEM (Gibco 11095-080) supplemented with 10% FBS (Gibco 26140-079), NEAA (Gibco 11140-050), GlutaMAX (Gibco 35050-061), Pen/Strep/Fungizone (Hyclone SV30079.01), and 2-mercaptoethanol (Gibco 21985-023). RPE-1 were cultured in DMEM/F12 (Gibco 11320-033) supplemented with 10% Fetal Plus (Atlas FP-0500-A) and Pen/Strep (Gibco 15140-122). For western blots, cells were grown in their respective media supplemented with 0.5% FBS starvation for 24 hours before harvesting to initiate ciliogenesis.

### Antisense oligonucleotides and transfections

Antisense oligonucleotides were purchased from IDT and fully modified with phosphorothioate linkages and 2′-methoxyethyl (MOE) modifications (**S. Table 1**). Cells were seeded 24 hours before transfecting ASOs using Lipofectamine 2000 (Invitrogen 11668027). 8 hours post-transfection, the media was replaced with fresh media. Note that the first ASO walk RT-PCR data comes from transfections where the media was not changed after 8 hours. For the western blots, cells were left to recover for 40 hours after replacing the media at 8 hours post-transfection. At 48 hours post-transfection, the media was replaced with 0.5% FBS starvation media. Cells were harvested 24 hours post-starvation.

### Genomic DNA isolation and allele phasing

Genomic DNA (gDNA) was isolated using AllPrep DNA/RNA Mini Kit (Qiagen 80204) and amplified using primers placed in exons 7 and 8. This reaction was used as the template for an additional round of PCR with primers in exon 7 and intron 7 (**S. Table 2**). KAPA HiFi HotStart PCR Kit (Roche 07958897001) was used with the cycling conditions: initial denaturation at 95°C for 3min, followed by 35 cycles of 98°C for 20s, 58°C for 15s, 72°C for 30s with a final extension of 72°C for 30s. Reactions were resolved on a 1% agarose gel and the product of interest was gel-purified using NucleoSpin Gel and PCR Clean-up kit (Macherey-Nagel 740609.250). The purified PCR product was A-tailed by incubating with Taq polymerase (NEB M0273) and 200μM dATPs (Thermo Scientific R0441) at 72°C for 15 minutes. Products were cloned into pCR 2.1 TOPO TA vector and transformed into NEB 5-alpha competent *E. coli* (NEB C2987H) using the TOPO TA Cloning Kit (Invitrogen 450641). Transformations were grown on kanamycin plates overnight and colonies were selected for overnight growth. Plasmid DNA was purified (Macherey-Nagel 740588.250) and sequenced (Quintara Biosciences).

### RNA isolation and cDNA synthesis

Cells were harvested and RNA was extracted using TRIzol Reagent (Invitrogen 15596026) and AllPrep DNA/RNA Mini Kit (Qiagen 80204) for allele phasing and first RT-PCR assay. All other RNA was isolated using RNeasy Micro Kit (Qiagen 74004). RNA was treated with DNase I Amplification grade (Invitrogen 18068015) and SuperScript III First-Strand Synthesis System was used to synthesize cDNA using random hexamers (Invitrogen 18080051).

### RNA sequencing

Poly-A selected total RNA was sequenced using NovaSeq X (80 million reads, paired end 2×150bp). Alignment was completed using the nf-core RNA-seq pipeline [14], including quality control, alignment, and visualization. Data was visualized using IGV_2.18.4. The analysis pipeline is available at https://github.com/rnabioco/jagannathan-aso-rnaseq-2024-06.

### RT-PCR

RT-PCR of *SDCCAG8* exons 7-8 for the ASO walk was performed using KAPA HiFi HotStart PCR Kit (Roche 07958897001) with the cycling conditions: 95°C for 3min, followed by 34 cycles of 98°C for 20s, 58°C for 15s, 72°C for 30s with a final extension of 72°C for 30s. Products were resolved on a 1% agarose gel. RT-PCR of SDCCAG8 exons 7-8 for the cryptic exon-targeting ASO walk, concentration course, amplicon sequencing was performed using the Phusion High-Fidelity DNA Polymerase (NEB M0530S) with the cycling conditions: 98°C for 3min, followed by 34 cycles of 98°C for 10s, 63°C for 15s, 72°C for 30s with a final extension of 72°C for 5min. Products were resolved on 2% agarose gels.

### Amplicon sequencing

Amplicons were purified from RT-PCR reactions using the NucleoSpin Gel and PCR Clean-up kit (Macherey-Nagel 740609.250). Big premium PCR Sequencing (6000 reads) was performed by Plasmidsaurus using Oxford Nanopore Technology. The R package dada2 was used to process raw fastq files and obtain unique amplicon sequences [15]. UCSC BLAT was used to confirm the identity of the unique amplicon sequences.

### Western blots

Protein samples were harvested in RIPA buffer (Thermo Fisher 89901) and quantified using Pierce BCA Protein assay kit (Thermo Fisher 23225). 30μg of protein was loaded on a 4– 12% Criterion XT Bis-Tris Protein Gel (Bio-Rad 3450123) in MOPS running buffer. Wet transfer was done using a nitrocellulose membrane in NuPAGE transfer buffer (Invitrogen NP00061).

Blots were blocked with Intercept (PBS) blocking buffer (LI-COR 927-70001) for 1 hour and incubated with rabbit anti-SDCCAG8 (1:500) (Novus Biologicals NBP2-13288) and mouse anti-GAPDH (1:1000) (Abcam ab9484) overnight at 4°C. Blots were then washed with PBS-T (0.1% Tween 20) (VWR EM8.22184.0500) 3 times with rocking for 10 min. Membranes were incubated with HRP secondary antibodies diluted 1:20,000 in blocking buffer with 0.2% Tween 20, for 1 hour with gentle rocking. Secondary antibodies used were: goat anti-rabbit (Thermo Fisher 31460) and goat anti-mouse (Thermo Fisher 31430). Blots were washed with PBS-T (0.05% Tween 20) with rocking for 5 min, this was repeated 5 times. The SuperSignal West Atto Ultimate Sensitivity Substrate (Thermo Scientific A38555) was used for protein detection.

## RESULTS

### Patient transcripts contain intronic regions between exons 7 and 8 and undergo nonsense-mediated decay (NMD)

To determine if the two mutations found in the patient were monoallelic or biallelic, genomic DNA was extracted from patient-derived fibroblasts and the region of intron 7 containing the mutations was amplified by PCR (**Fig. 1a**). These amplicons were cloned into TOPO vectors and sequenced; each plasmid contained only one of the two mutations, indicating that the mutations were biallelic (**Fig. 1b**).

**Figure 1.**
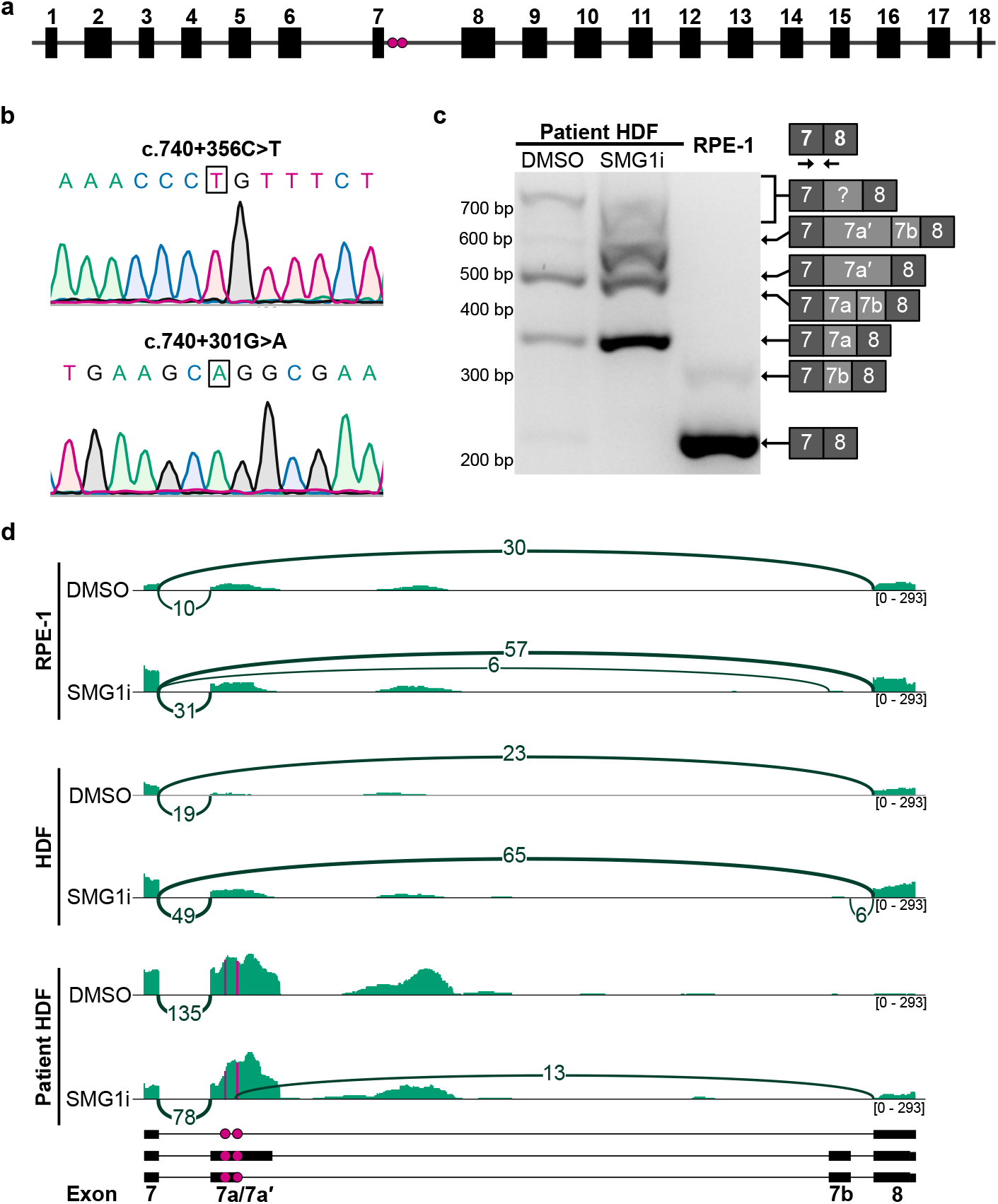
Characterizing patient mutations by allele phasing, splicing assay, and RNA sequencing. a) Canonical *SDCCAG8* gene diagram (ENST00000366541.8). Magenta dots indicate the mutations, c.740+301G>A (left) and c.740+356C>T (right). b) Sequencing traces for patient allele phasing. c) RT-PCR assay to assess splicing between exons 7 and 8 in patient cells with primers flanking intron 7 (left), diagram of amplicons (right); Expected size of canonical splicing product (exon 7-8) is 222bp. d) Sashimi plot of paired-end RNA-seq (40M read pairs) of patient cells and unaffected cell lines +/-SMG1i (NMD inhibitor). Minimum splice junction threshold set at 4. Mutations marked with magenta lines and dots, c.740+301G>A (left) and c.740+356C>T (right). Tracks were autoscaled by group using IGV.

To assess the effect of both mutations on splicing, an RT-PCR assay was performed using primers targeting exons 7 and 8. In patient-derived fibroblasts, there were multiple PCR products larger than those produced from unaffected RPE-1 cells, suggesting the inclusion of intronic regions (**Fig. 1c**). Patient cells were treated with SMG1i, a small molecule inhibitor of the SMG1 kinase involved in NMD [16], which resulted in the stabilization of mis-spliced transcripts as observed by RT-PCR and RNA-seq (**Fig. 1c-d**, and **Fig. S1**). RT-PCR amplicons were sequenced, confirming the inclusion of exons 7a (112bp) and 7b (97bp), both of which are known to be associated with c.740+356C>T [7]. We also observed exon 7a′ (277bp), which shares a 3′ splice site with 7a (**Fig. S1**). RNA sequencing of these samples confirmed a complete absence of normal splicing from exons 7 to 8 in patient cells (**Fig. 1d**). Based on these data, we concluded that mutations found in the BBS patient alter the splicing of *SDCCAG8* transcripts and could likely be treated with splice-switching ASOs to prevent the inclusion of cryptic exons.

### ASO walk identifies oligonucleotides that alter splicing and increase the level of full-length *SDCCAG8*transcript

To restore normal splicing of *SDCCAG8* transcripts in the patient-derived fibroblasts, ASOs were tiled over each mutation to identify splice-switching candidates. 20mer 2′MOE PS ASOs were shifted by 2nt over each mutation, with ASOs 1-9 targeting the known mutation c.740+356C>T and ASOs 10-18 targeting the VUS c.740+301G>A (**Fig. 2a**). Patient-derived fibroblasts were transfected with 100nM ASOs, and total RNA was harvested after 24 hours. RT-PCR was performed to identify ASOs that increased canonical splicing between exons 7 and 8 (**Fig. 2b**). ASOs 2-4, 6, and 12 displayed an increase in the canonical splicing product, with ASOs 4 and 12 being chosen for further investigation based on the intensity of their bands.

**Figure 2.**
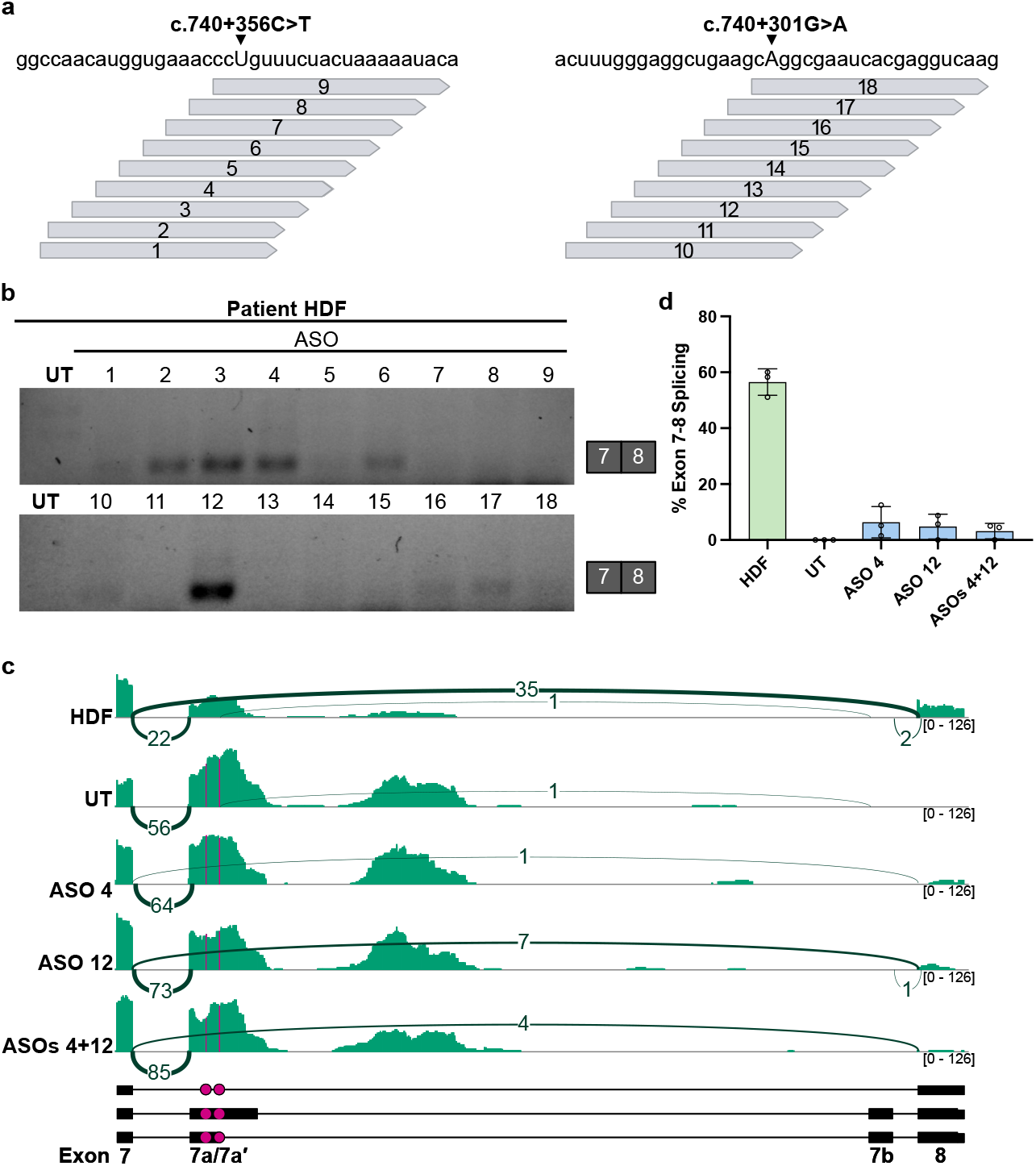
ASO-mediated *SDCCAG8* mRNA splicing between exons 7 and 8. a) Diagram of ASOs tiled over mutations (5′-3′). Mutations indicated by triangles and capital letters. b) RT-PCR of patient cells treated with ASOs; Expected size of canonical splicing product (exon 7-8) is 222bp. n=1. c) Sashimi diagram of patient cells treated with ASOs. Tracks were autoscaled by group using IGV and all splice junctions are shown. Mutation locations indicated with magenta lines and dots. Representative sashimi plots chosen out of 3 biological replicates of each. d) Percent of exon 7 to 8 splice junctions out of total splice junctions for RNA-seq replicates.

To further evaluate the effect of ASOs 4 and 12 on splicing, poly-A RNA sequencing was performed for patient-derived fibroblasts transfected with ASO 4, ASO 12, and a dual treatment (ASOs 4+12). An increase in read coverage over the splice junction between exons 7 and 8 was observed along with increased exonic coverage downstream of the mutations, indicating successful exclusion of cryptic exons (**Fig. 2c; Fig. S2**). The average ratio of splicing from exon 7 to 8 increased from 0% in untreated patient cells to 6.4%, 4.8%, and 3.2% for ASOs 4, 12, and 4+12, respectively (**Fig. 2d**). The observed changes in splicing suggest reduced NMD of *SDCCAG8* transcripts due to ASO-induced exclusion of cryptic exon(s).

We then directly targeted the mis-splicing by walking ASOs over the 3′ splice sites of the cryptic exons 7a/7a′ and 7b. These ASOs were then narrowed down to 3 candidate ASOs per 3′ splice site by the number of off-target binding sites found using UCSC BLAT (**Fig. 3a**). Patient-derived fibroblasts were transfected with each ASO at 100nM. RNA was harvested and RT-PCR was performed to assess splicing. ASOs 19-21, targeting exon 7a/7a′, each increased splicing from exon 7 to 8 (**Fig. 3b**). ASOs 22-24 all appeared to stabilize a ∼500bp product but did not increase the 222bp product that would indicate canonical splicing. ASO 20 was chosen for further investigation because its GC% was between 40-60% and it did not produce significant aberrant products below 200bp like ASO 21, suggesting more specific splice-switching activity. RNA-seq of ASO 20 treated cells confirmed increased canonical splicing compared to the untreated patient cells (**Fig. 3c; Fig S3**). The ratio of splicing from exon 7 to 8 increased from 0% in untreated cells to an average of 26% with ASO 20 treatment (**Fig. 3d**), representing a 4-5 fold improvement over the mutation-targeting ASOs 4 and 12 (**Fig. 2**).

**Figure 3.**
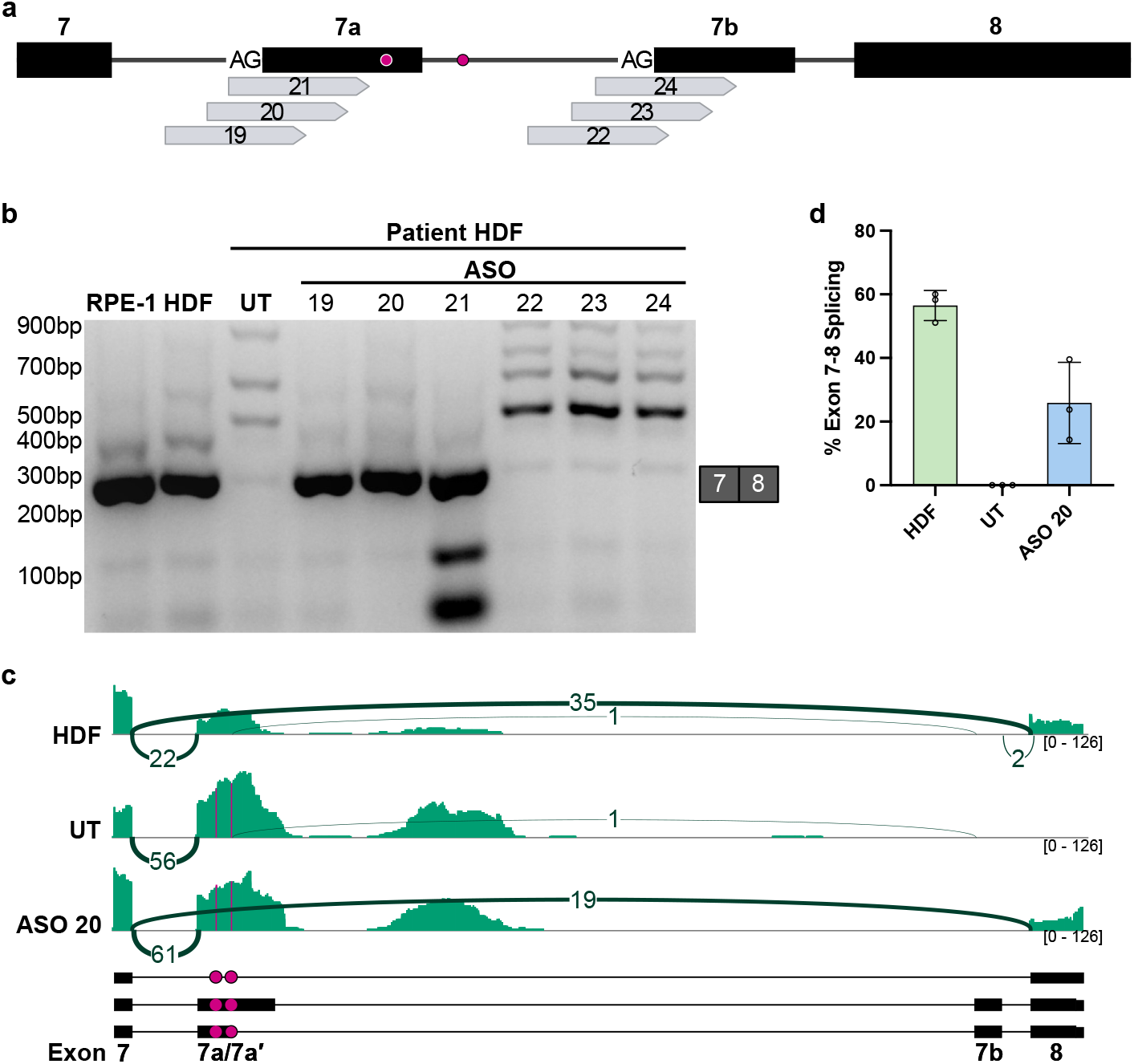
ASOs targeting cryptic exons alter *SDCCAG8* mRNA splicing between exons 7 and 8. a) Diagram of ASOs targeting exons 7a and 7b (not to scale). Magenta dots indicate the mutations, c.740+301G>A (left) and c.740+356C>T (right). b) RT-PCR of patient cells treated with 100nM ASOs 19-24. Expected size of canonical splicing product (exon 7-8) is 222bp. c) Sashimi plot of patient cells treated with ASO 20. Tracks were autoscaled by group using IGV and all splice junctions are shown. (Same HDF and UT tracks as Fig. 2). d) Percent of exon 7 to 8 splice junctions out of total splice junctions for RNA-seq replicates.

#### *SDCCAG8* splicing product and protein levels increase with ASO treatment

To determine if the normal *SDCCAG8* transcript increases with increasing ASO concentration, patient-derived fibroblasts were transfected with 4, 20, and 100nM ASOs, and RT-PCR was performed (**Fig. 4a-b**). The intensity of the wild-type bands increased from 4nM to 20nM for each ASO condition, but slightly decreased or plateaued from 20nM to 100nM, with the exception of ASO 20, which continued to show increased splicing at 100nM. Each ASO condition was significantly different from untreated patient-derived fibroblasts regardless of concentration, demonstrating robust and reproducible splice correction across a range of ASO doses (**Fig. 4b**).

**Figure 4.**
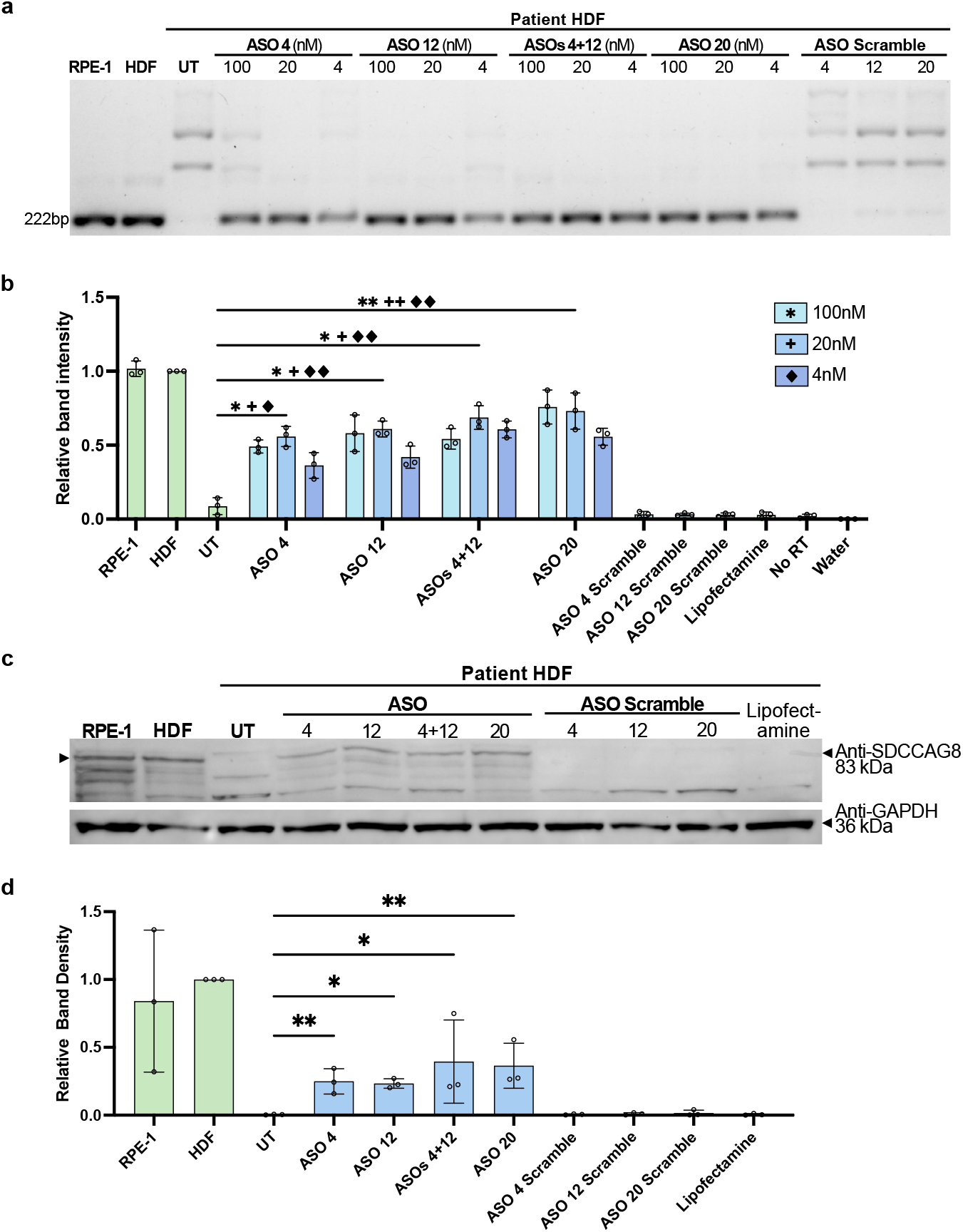
Impact of ASO treatment on *SDCCAG8* levels. a) RT-PCR of patient cells treated with 4nM, 20nM, or 100nM ASOs for 8hours and harvested 24hours after treatment. b) Quantification of the 222bp band in (a) and biological replicates (not shown). Values normalized to the corresponding HDF band. Asterisks represent significance between 100nM ASOs, plus signs for 20nM, and diamonds for 4nM. c) Western blot of patient cells transfected with ASOs and serum starved for 24hours before harvest. Blotted for SDCCAG8 (83 kDa) and GAPDH (36 kDa). d) Quantification of band intensity in (c) and biological replicates. Values normalized to GAPDH and the corresponding HDF condition. (b and d) Significance calculated using RM-one way ANOVA with the Geisser-Greenhouse correction and an Uncorrected Fisher’s LSD with individual variances computed for each comparison. (P ≤ 0.05 (*), ≤ 0.01 (**)).

To investigate if the exclusion of cryptic exon(s) in ASO-treated patient-derived fibroblasts causes an increase in SDCCAG8 protein, patient cells were transfected with 100nM ASOs and serum-starved to induce ciliogenesis. Western blots were performed with three biological replicates and revealed significant increases in SDCCAG8 protein (**Fig. 4c-d**). After normalization to GAPDH and HDF bands, SDCCAG8 protein significantly increased in ASO-treated patient-derived fibroblasts, consistent with the exclusion of PTC-containing cryptic exons that would trigger NMD (**Fig. 4a**). Importantly, the restored protein migrated at the expected molecular weight of 83 kDa, confirming production of full-length SDCCAG8 containing the functionally critical C-terminal domain. Thus, increasing the concentration of ASO treatment led to increased canonical splicing from exons 7 to 8 in *SDCCAG8* transcripts. Recovery of canonical splicing by ASO treatment increased SDCCAG8 protein from undetectable levels in untreated patient cells to approximately 20-40% of the levels observed in control fibroblasts, with ASO 20 showing the largest, most significant protein restoration across conditions.

## DISCUSSION

Patient-derived mutations cause the inclusion of cryptic exons between *SDCCAG8* exons 7 and 8 that disrupt the reading frame and result in the degradation of the transcripts by NMD. Although these mutations induce alternative splicing, the exact mechanism is unknown. The mutation c.740+356C>T is predicted to promote alternative splicing through one of two mechanisms: the first is the loss of an exonic splicing enhancer binding site, and the second is through Alu element-mediated alternative splicing [7,17]. Regardless of the mechanism used to include these cryptic exon(s) in the mRNA, splicing can be altered by ASOs targeting sequence elements recognized by the spliceosome.

When these mutations were targeted with ASOs, we observed increased canonical splicing in patient-derived fibroblasts. We found that ASOs 4, 12, 4+12, and 20 increase canonical splicing of the *SDCCAG8* transcript between exons 7 and 8. The amount of canonical splicing increased with ASO concentration from 4nM to 20nM. The effect of the ASOs may plateau at 20nM, as the trend of increased canonical splicing did not continue at 100nM. Out of all ASO conditions, ASO 20 had the greatest effect at 100nM and 20nM, while the combination of ASOs 4 and 12 had the greatest effect at 4nM. ASO 20 may outperform the other ASOs because it blocks access to the 3′ splice site of exon 7a/7a′ rather than blocking unknown upstream events that may occur due to the mutations. When the ratio of splicing directly from exons 7-8 is calculated from the sashimi plots, ASO 20 has the largest average ratio of 26% while ASOs 4,12, and 4+12 have ratios of 6.4%, 4.8%, and 3.2% (**Fig. 2d; S2, 3d; S3**). Although these percentages are lower than the 57% splicing observed in control fibroblasts, ASO 20 recovers nearly half of this splicing from the 0% seen in untreated patient cells. When the PTC-containing cryptic exon(s) are excluded from the final transcript, NMD does not degrade the *SDCCAG8* transcripts, which should result in increased protein output. Consistent with this expectation, ASO treatment restored full-length SDCCAG8 protein from undetectable levels to approximately 20-40% of wild-type expression. While partial, this level of restoration may be therapeutically meaningful, as BBS is a recessive disorder requiring complete loss of SDCCAG8 for disease manifestation, and precedent from other genetic diseases demonstrate that partial protein restoration can provide clinical benefit [18].

Splice-altering ASOs prevent the spliceosome from recognizing aberrant signals that induce splicing of PTC-containing cryptic exons. ASOs targeting this patient’s mutations and the cryptic exon splice site are expected to restore the C-terminus of SDCCAG8 and its ability to localize to the centrosome. SDCCAG8 contributes to vesicle docking to the basal body and promotes cilia formation and function [7,8,19]. For example, the C-terminus of SDCCAG8 interacts with ICK and MAK and a restoration of these interactions may allow these proteins to promote ciliogenesis and regulate cilia length [12,20].

The primary goal of ASO treatment for this patient is to slow or halt BBS-related retinal degeneration. Photoreceptor outer segments are specialized primary cilia that share many of the same proteins and ciliary assembly mechanisms as canonical primary cilia [21]. Restoration of SDCCAG8 expression in photoreceptors could therefore ameliorate progressive retinal degeneration. Administering ASOs to this patient could treat their retinal degeneration by restoring SDCCAG8 expression.

While the c.740+301G>A substitution has not been previously reported in BBS patients, c.740+356C>T has been previously associated with cryptic exon insertion in *SDCCAG8*. Exon 7b has also been observed at a low frequency in control cell lines such as HDF and RPE-1. If the mutations observed in this patient increase Alu-mediated exon inclusion, others who have mutations in the same region of the gene may share the same cryptic exon insertions [17]. Mutation-agnostic ASOs such as ASO 20 can treat any patient with these insertions. Identifying splicing events that increase in frequency when there are mutations present could help to design ASOs to treat a larger patient population than designing ASOs for individual mutations.

While ASO treatment restores protein expression in patient-derived fibroblasts, this model is not representative of photoreceptors or other cell types that would receive ASOs during treatment. Despite this limitation, testing therapeutics in patient-derived fibroblasts allows efficient and rapid testing of different conditions in a model that contains mutations of interest. Patient-derived fibroblasts are easier to obtain than other patient-derived models that would be more relevant to the eye, such as retinal organoids. Another limitation of these experiments is the use of transfection reagents, which can cause cell stress, and the lack of physiological relevance. While we demonstrate robust protein-level rescue meeting field standards for ASO preclinical validation, future studies in more disease-relevant models will assess functional outcomes, including ciliogenesis and photoreceptor function.

Although treatments for the symptoms of BBS are available to patients, currently there is no targeted therapy for the genetic causes of this patient’s retinal degeneration. ASO treatments can recover SDCCAG8 protein expression in patient-derived fibroblasts, which could help restore the function of this protein in photoreceptors. Clinical translation of the lead ASOs we have identified in this study will require chemistry optimization to compare 2′MOE phosphorothioate ASOs with alternative chemistries, *in vivo* efficacy studies in animal models to assess functional rescue, delivery optimization for intravitreal or subretinal injection, and standard IND-enabling toxicology studies. Given the progressive retinal degeneration in this patient, expedited development pathways similar to those used for other N-of-1 antisense therapies may be applicable [4].

## CONCLUSIONS

We developed and validated antisense oligonucleotides that correct mutation-induced aberrant splicing in *SDCCAG8*, restoring full-length protein from undetectable levels to approximately 20-40% of wild-type in patient-derived fibroblasts. ASO 20, which targets the cryptic exon splice site rather than patient-specific mutations, demonstrated superior efficacy and mutation-agnostic therapeutic potential. These ASOs add to the rising number of therapies for rare diseases that previously had no targeted treatment. While these kinds of therapeutics open new avenues to improving patient care beyond current symptom-focused treatments, it is difficult to develop a single ASO that can treat multiple patients. ASO 20’s mutation-agnostic mechanism could benefit any patient with cryptic exon inclusion in this region of *SDCCAG8*, addressing this challenge. Identifying aberrant splicing that occurs at low frequencies in unaffected cells could facilitate the identification of new therapeutic targets shared between larger populations of patients than ASOs typically treat. This work demonstrates the therapeutic potential of splice-switching ASOs for ciliopathies and highlights the value of targeting common downstream splicing defects to maximize patient benefit.

## ACKNOWLEDGEMENTS

We thank Matthew Taliaferro for the suggestion of targeting cryptic exon splicing independently of mutation-specific targeting. We thank the Gates Institute Stem Cell Biobank and Disease Modeling Core for deriving fibroblasts from the patient’s skin punch biopsy. We thank members of the Jagannathan and Hesselberth laboratory for their insightful feedback on the manuscript. This work was supported by the RNA Bioscience Initiative (K.E.M.)., Children’s Hospital of Colorado (K.E.M. and A.E.C.), R35 GM119550 (J.R.H.), University of Colorado School of Medicine Translational Research Scholars Program (S.J.), and the National Institutes of Health grant R35GM133433 (S.J.).

## DECLARATION OF GENERATIVE AI AND AI-ASSISTED TECHNOLOGIES

During the preparation of this work, the authors used Claude.ai to improve language and readability. After using this tool, the authors reviewed and edited the content as needed and take full responsibility for the content of the publication.

## SUPPLEMENTAL INFORMATION

**Supplementary Table 1:**
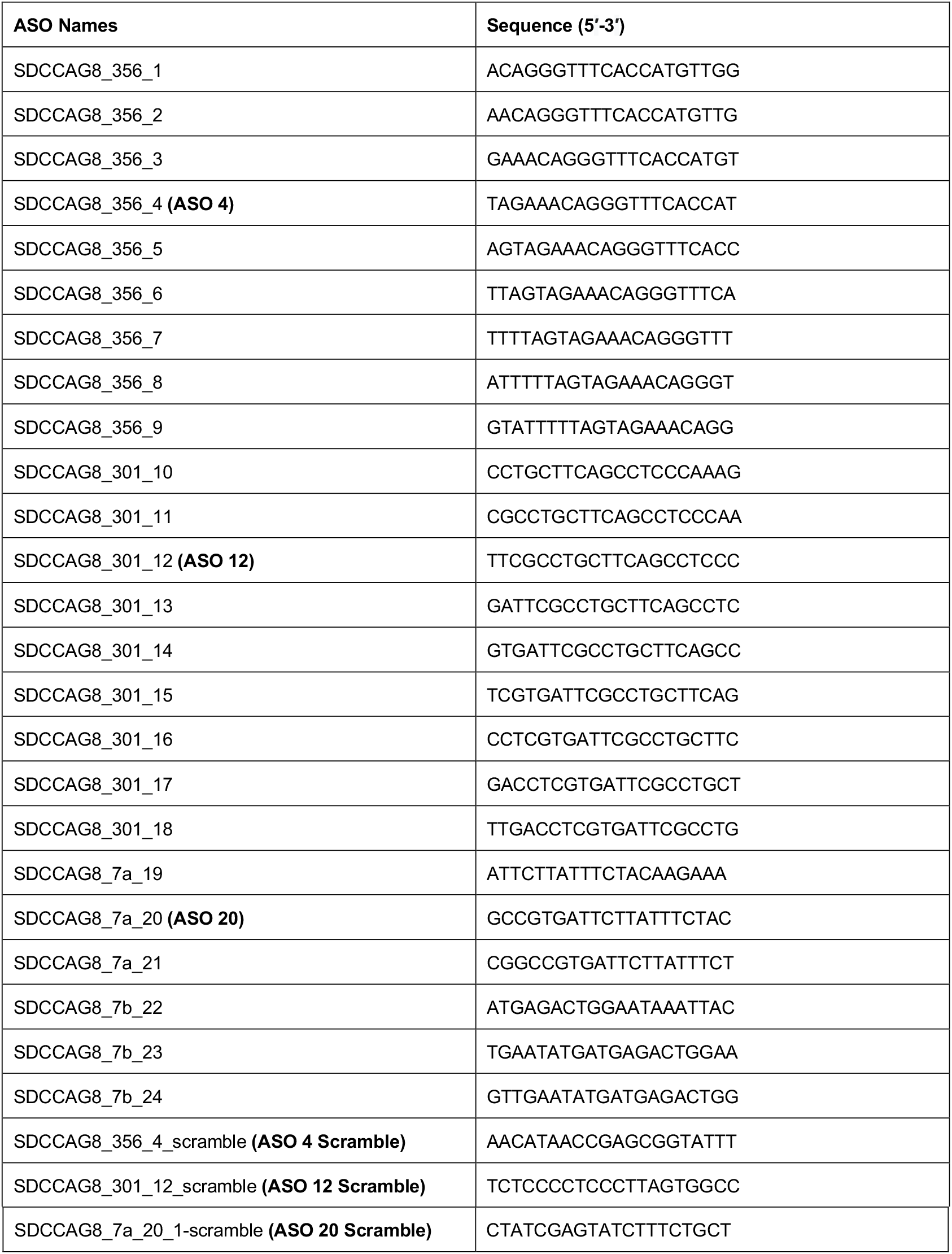

**Supplementary Table 2:**
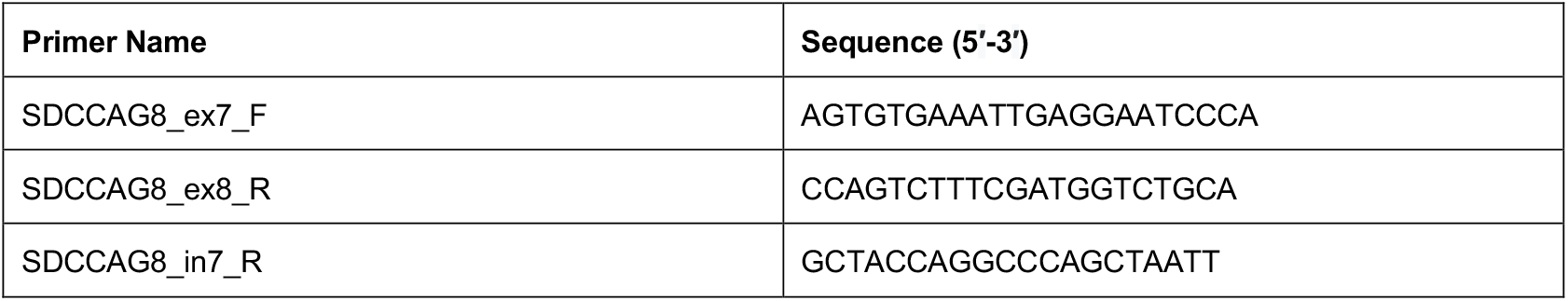

## SUPPLEMENTARY FIGURES

**Supplementary Figure 1.**
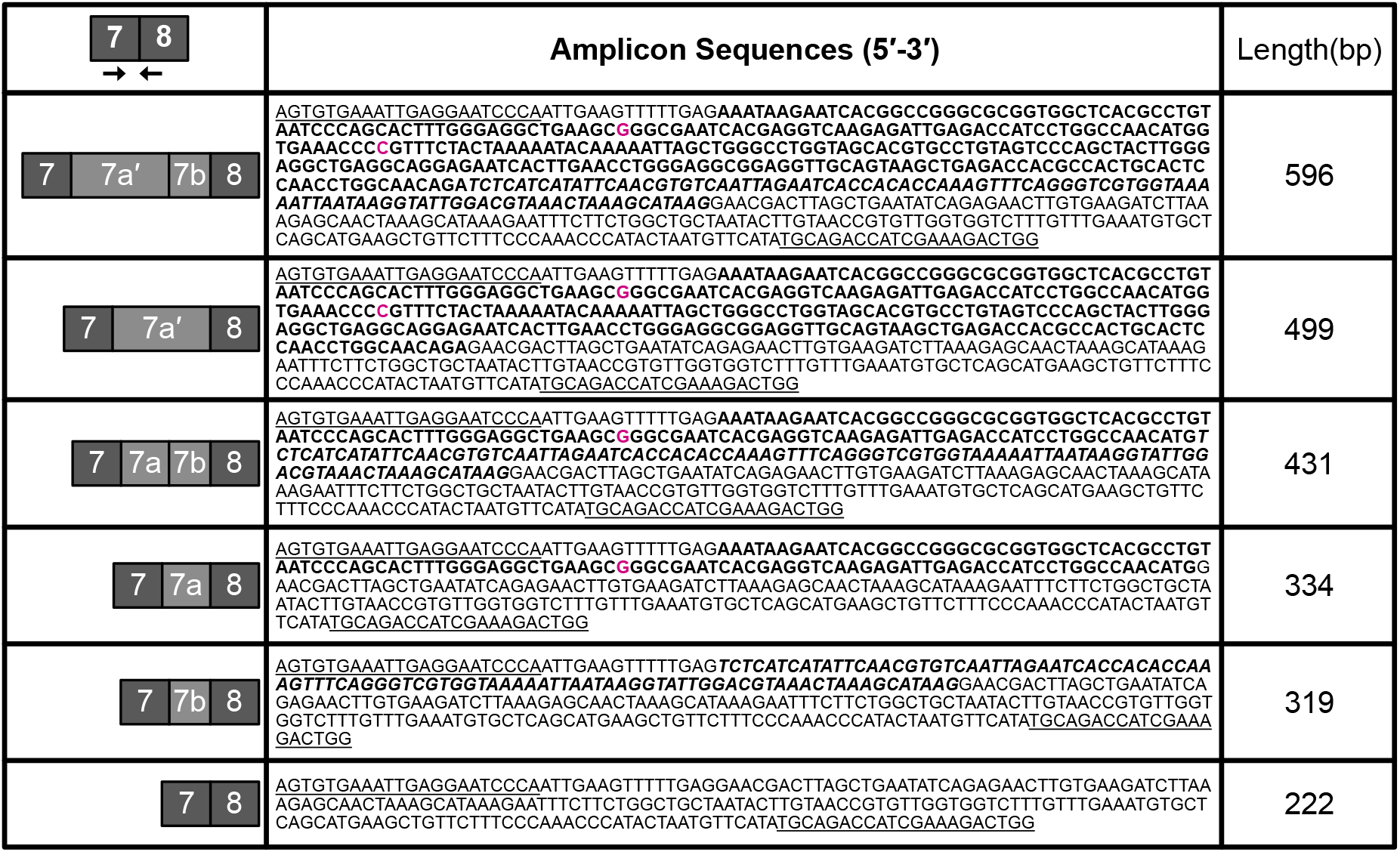
Amplicon Sequences obtained from splicing assay. PCR reactions using primers in SDCCAG8 exons 7 and 8 and cDNA from RPE-1, HDF (Sigma 106-05A) and patient-derived fibroblasts were purified over columns. Purified reactions were sent for amplification-free long read Oxford Nanopore sequencing. Representative splicing diagrams are shown next to amplicon sequences and lengths. Annotations: primer sequences are underlined, exons 7a and 7a′ are bold, exon 7b is bold and italicized, locations of mutations are in magenta. Both mutations, c.740+301G>A and c.740+354C>T, were observed separately in the amplicons containing mutation locations.

**Supplementary Figure 2.**
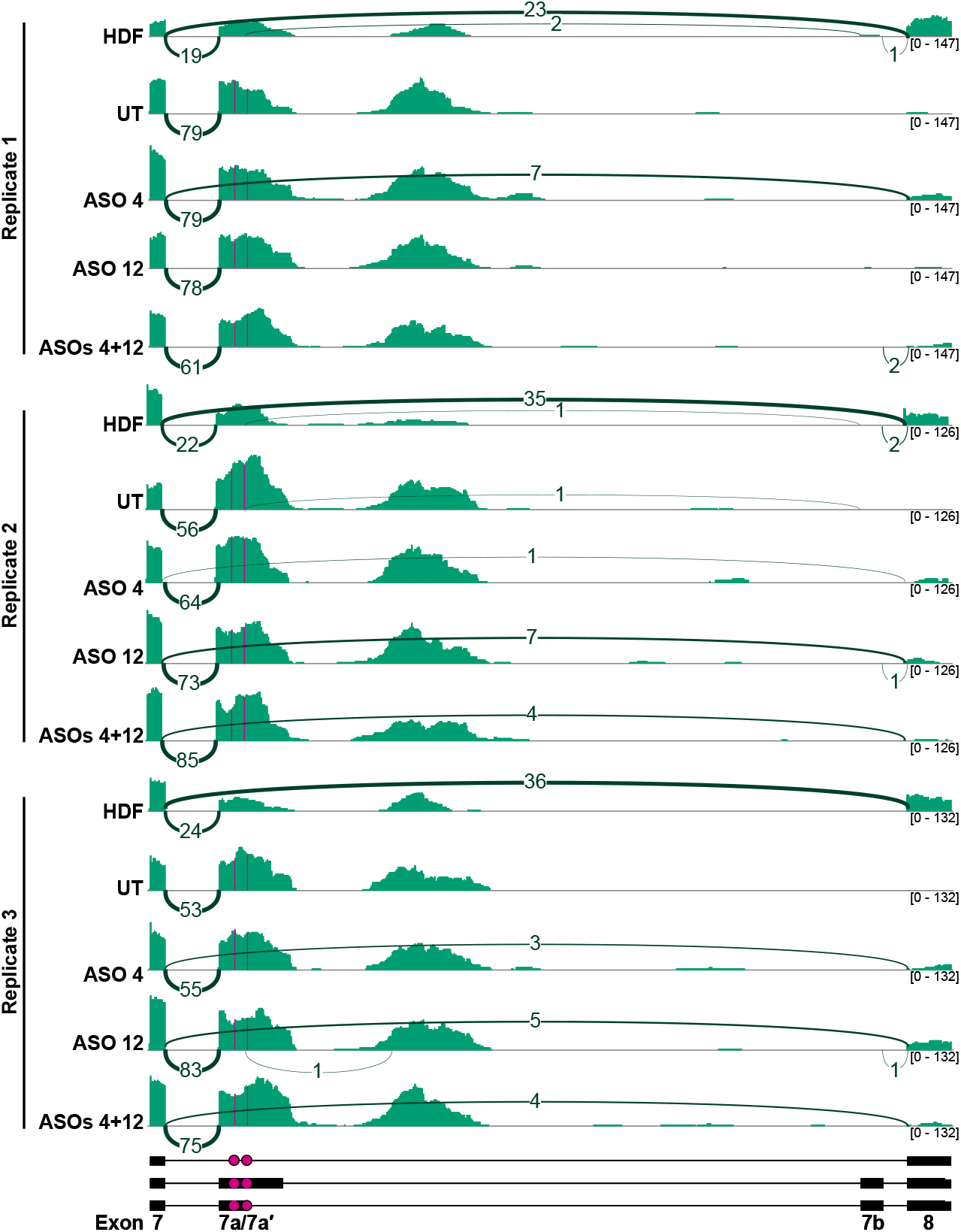
RNA-seq sashimi diagram of patient cells treated with ASOs. Tracks were autoscaled by group using IGV and all splice junctions are shown. Mutation locations indicated with magenta lines and dots.

**Supplementary Figure 3.**
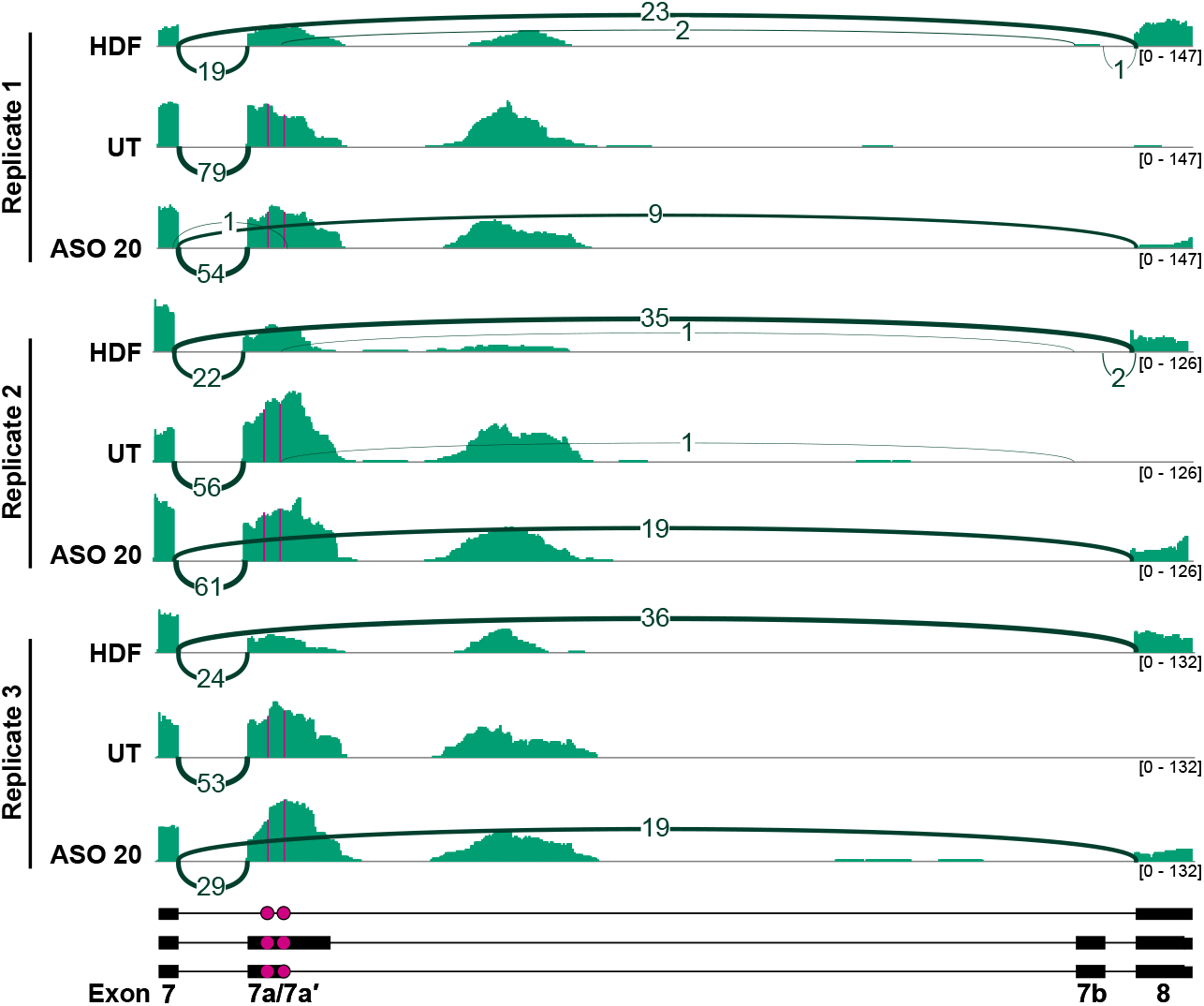
RNA-seq sashimi diagram of patient cells treated with ASOs. Tracks were autoscaled by group using IGV and all splice junctions are shown. (Same HDF and UT tracks as Fig. S2). Mutation locations indicated with magenta lines and dots.

## REFERENCES

1. Han Z, Chen C, Christiansen A, Ji S, Lin Q, Anumonwo C, et al. Antisense oligonucleotides increase Scn1a expression and reduce seizures and SUDEP incidence in a mouse model of Dravet syndrome. Sci Transl Med. 2020;12:eaaz6100. 10.1126/scitranslmed.aaz6100

2. Singh NN, Lee BM, DiDonato CJ, Singh RN. Mechanistic principles of antisense targets for the treatment of Spinal Muscular Atrophy. Future Med Chem. 2015;7:1793–808. 10.4155/fmc.15.101

3. Aartsma-Rus A, Garanto A, van Roon-Mom W, McConnell EM, Suslovitch V, Yan WX, et al. Consensus Guidelines for the Design and In Vitro Preclinical Efficacy Testing N-of-1 Exon Skipping Antisense Oligonucleotides. Nucleic Acid Ther. 2023;33:17–25. 10.1089/nat.2022.0060

4. Kim J, Hu C, Achkar CME, Black LE, Douville J, Larson A, et al. Patient-Customized Oligonucleotide Therapy for a Rare Genetic Disease. New England Journal of Medicine. Massachusetts Medical Society; 2019;381:1644–52. 10.1056/NEJMoa1813279

5. Melluso A, Secondulfo F, Capolongo G, Capasso G, Zacchia M. Bardet-Biedl Syndrome: Current Perspectives and Clinical Outlook. Ther Clin Risk Manag. 2023;19:115–32. 10.2147/TCRM.S338653

6. Kenedy AA, Cohen KJ, Loveys DA, Kato GJ, Dang CV. Identification and characterization of the novel centrosome-associated protein CCCAP. Gene. 2003;303:35–46. 10.1016/S0378-1119(02)01141-1

7. Otto EA, Hurd TW, Airik R, Chaki M, Zhou W, Stoetzel C, et al. Candidate exome capture identifies mutation of SDCCAG8 as the cause of a retinal-renal ciliopathy. Nat Genet. 2010;42:840–50. 10.1038/ng.662

8. Airik R, Schueler M, Airik M, Cho J, Ulanowicz KA, Porath JD, et al. SDCCAG8 Interacts with RAB Effector Proteins RABEP2 and ERC1 and Is Required for Hedgehog Signaling. PLoS One. 2016;11:e0156081. 10.1371/journal.pone.0156081

9. Flynn M, Whitton L, Donohoe G, Morrison CG, Morris DW. Altered gene regulation as a candidate mechanism by which ciliopathy gene SDCCAG8 contributes to schizophrenia and cognitive function. Human Molecular Genetics. 2020;29:407–17. 10.1093/hmg/ddz292

10. Submissions for variant NM_006642.5(SDCCAG8):c.740+301G>A - ClinVar Miner [Internet]. [cited 2024 Nov 25]. https://clinvarminer.genetics.utah.edu/submissions-by-variant/NM_006642.5%28SDCCAG8%29%3Ac.740%2B301G%3EA. Accessed 25 Nov 2024

11. Dyle MC, Kolakada D, Cortazar MA, Jagannathan S. How to get away with nonsense: Mechanisms and consequences of escape from nonsense-mediated RNA decay. WIREs RNA. 2020;11:e1560. 10.1002/wrna.1560

12. Tsutsumi R, Chaya T, Tsujii T, Furukawa T. The carboxyl-terminal region of SDCCAG8 comprises a functional module essential for cilia formation as well as organ development and homeostasis. Journal of Biological Chemistry. 2022;298:101686. 10.1016/j.jbc.2022.101686

13. Li K, Zhou X, Liu W, Wang Y, Zhang Z, Zhang H, et al. Loss of C-Terminal Coiled-Coil Domains in SDCCAG8 Impairs Centriolar Satellites and Causes Defective Sperm Flagellum Biogenesis and Male Fertility. Cells. 2025;14:1135. 10.3390/cells14151135

14. Ewels PA, Peltzer A, Fillinger S, Patel H, Alneberg J, Wilm A, et al. The nf-core framework for community-curated bioinformatics pipelines. Nat Biotechnol. Nature Publishing Group; 2020;38:276–8. 10.1038/s41587-020-0439-x

15. Callahan BJ, McMurdie PJ, Rosen MJ, Han AW, Johnson AJA, Holmes SP. DADA2: High-resolution sample inference from Illumina amplicon data. Nat Methods. Nature Publishing Group; 2016;13:581–3. 10.1038/nmeth.3869

16. Gopalsamy A, Bennett EM, Shi M, Zhang W-G, Bard J, Yu K. Identification of pyrimidine derivatives as hSMG-1 inhibitors. Bioorganic & Medicinal Chemistry Letters. 2012;22:6636–41. 10.1016/j.bmcl.2012.08.107

17. Borovská I, Vořechovský I, Královičová J. Alu RNA fold links splicing with signal recognition particle proteins. Nucleic Acids Research. 2023;51:8199–216. 10.1093/nar/gkad500

18. Mittal S, Tang I, Gleeson JG. Evaluating human mutation databases for “treatability” using patient-customized therapy. Med. 2022;3:740–59. 10.1016/j.medj.2022.08.006

19. Senatore E, Iannucci R, Chiuso F, Delle Donne R, Rinaldi L, Feliciello A. Pathophysiology of Primary Cilia: Signaling and Proteostasis Regulation. Front Cell Dev Biol [Internet]. Frontiers; 2022 [cited 2025 Jan 10];10. 10.3389/fcell.2022.833086

20. Chaya T, Maeda Y, Tsutsumi R, Ando M, Ma Y, Kajimura N, et al. Ccrk-Mak/Ick signaling is a ciliary transport regulator essential for retinal photoreceptor survival. Life Sci Alliance. 2024;7:e202402880. 10.26508/lsa.202402880

21. Chen HY, Kelley RA, Li T, Swaroop A. Primary cilia biogenesis and associated retinal ciliopathies. Semin Cell Dev Biol. 2021;110:70–88. 10.1016/j.semcdb.2020.07.013

